# GeneKnow: An auditable AI framework for source-grounded biological evidence synthesis

**DOI:** 10.64898/2026.05.28.728511

**Authors:** Hongpan Zhang, Bhummanat Sittipongpittaya, Chongzhi Zang

**Affiliations:** Department of Genome Sciences, University of Virginia, Charlottesville, VA, USA; Department of Biochemistry and Molecular Genetics, University of Virginia, Charlottesville, VA, USA; Department of Biomedical Engineering, University of Virginia, Charlottesville, VA, USA; UVA Comprehensive Cancer Center, Charlottesville, VA, USA

## Abstract

Biological function is context dependent, yet synthesizing evidence for a gene’s function in a defined cell type, disease, or perturbation remains labor-intensive. Here we present GeneKnow, an auditable artificial intelligence (AI) framework for source-grounded biological evidence synthesis. GeneKnow separates evidence-critical operations, including literature retrieval, passage selection, provenance tracking, and bibliography construction, from generative steps that are constrained to semantic analysis and literature synthesis. GeneKnow supports multi-paper discovery and single-paper inspection while preserving links from synthesized claims to source passages, generating trustworthy syntheses without fabricated citations and minimizing hallucinations. Systematic benchmarking showed that GeneKnow achieved higher claim support and citation fidelity than leading general-purpose and scientific AI systems. These results demonstrate that an AI system with controlled division of labor between deterministic and generative components can substantially improve the fidelity and auditability of biomedical literature synthesis.

## Introduction

Understanding context-dependent gene function is a fundamental goal of molecular biology, because the same gene can exert distinct or even opposing effects across cell types, disease states, developmental stages, and experimental perturbations^1–3^. Determining the role of an individual gene in a defined context therefore requires integrating mechanistic evidence distributed across multiple studies. Gene set enrichment and functional annotation methods efficiently summarize pathways associated with gene lists, but they offer limited insight into the roles of individual genes in a specific context^4–6^. Researchers therefore often rely on labor-intensive manual literature review to interpret gene-context relationships. The same burden also appears during computational method development, where published studies are frequently used as ground-truth evidence for training, validating, or benchmarking machine-learning models, which further increases the need for efficient and reliable literature synthesis.

Large language models (LLMs) create new opportunities to automate scientific literature synthesis, but their use in research is still constrained by factual reliability, citation integrity, and auditability. General-purpose LLMs are prone to hallucination, generating plausible but unsupported claims and fabricated references^7,8^. Retrieval-augmented generation (RAG) can reduce unsupported generation by conditioning models on retrieved evidence, although some implementations, including OpenScholar^9^, depend on large pre-indexed corpora or local databases that require substantial storage and maintenance. Passage-level vector search may retrieve locally relevant text fragments but miss related evidence within the same paper, limiting its ability to support complete and context-aware evidence extraction. Moreover, it may lack reproducibility due to non-deterministic indexing and approximate searches^10,11^. Web-enabled research agents provide broader access to current information, but their search depth, retrieval behavior, and citation workflows can vary across runs and are often difficult to audit^12,13^. A scientific synthesis system must therefore do more than opaque evidence selection and fluent prose generation; it should make evidence retrieval, citation construction, and claim generation auditable and reproducible throughout the literature synthesis process.

Recent systems have advanced scientific literature synthesis through adaptive retrieval, citation-aware reasoning^14^, and structured integration of biological databases^15^. However, most are optimized for open-ended exploration or factual database queries rather than for reconstructing context-specific gene functions from primary literature. We hypothesized that scientific reliability could be improved through an explicit division of labor between deterministic and generative computation. Operations for which exactness and reproducibility are essential, including literature retrieval, evidence selection, provenance tracking, and citation construction, should remain deterministic, whereas generative models should be reserved for semantically demanding but evidence-constrained tasks such as summarization, claim verification, and synthesis.

To test this principle, we developed GeneKnow, an auditable framework for source-grounded gene-context evidence synthesis. GeneKnow Discover retrieves and integrates evidence across multiple relevant papers, whereas GeneKnow Inspect extracts and summarizes evidence from a designated study. We evaluated the contributions of evidence-focused hierarchical summarization and claim-level self-verification through ablation experiments, and benchmarked end-to-end performance against leading general-purpose and literature-focused AI systems, including ChatGPT 5.2 Thinking, Claude Opus 4.6, Gemini 3 Thinking, and OpenScholar. Our results show that this controlled architecture achieves high claim support and citation fidelity without sacrificing evidence yield, providing a broadly applicable design strategy for auditable AI-assisted biomedical synthesis.

## Results

### A controlled division of labor enables auditable biological evidence synthesis

We designed GeneKnow around an explicit division of labor between deterministic evidence operations and constrained generative interpretation (Fig. 1). Rather than asking a single model to retrieve, interpret, cite, and synthesize literature in one unconstrained process, GeneKnow assigns paper retrieval, evidence selection, provenance tracking, and bibliography construction to deterministic procedures. Generative models are used only for tasks requiring semantic interpretation, including passage summarization, article-level integration, claim verification, and cross-paper synthesis. By default, GeneKnow performs these generative operations through the OpenAI API using gpt-5-mini, while its modular architecture permits substitution of other LLM backends.

**Figure 1.**
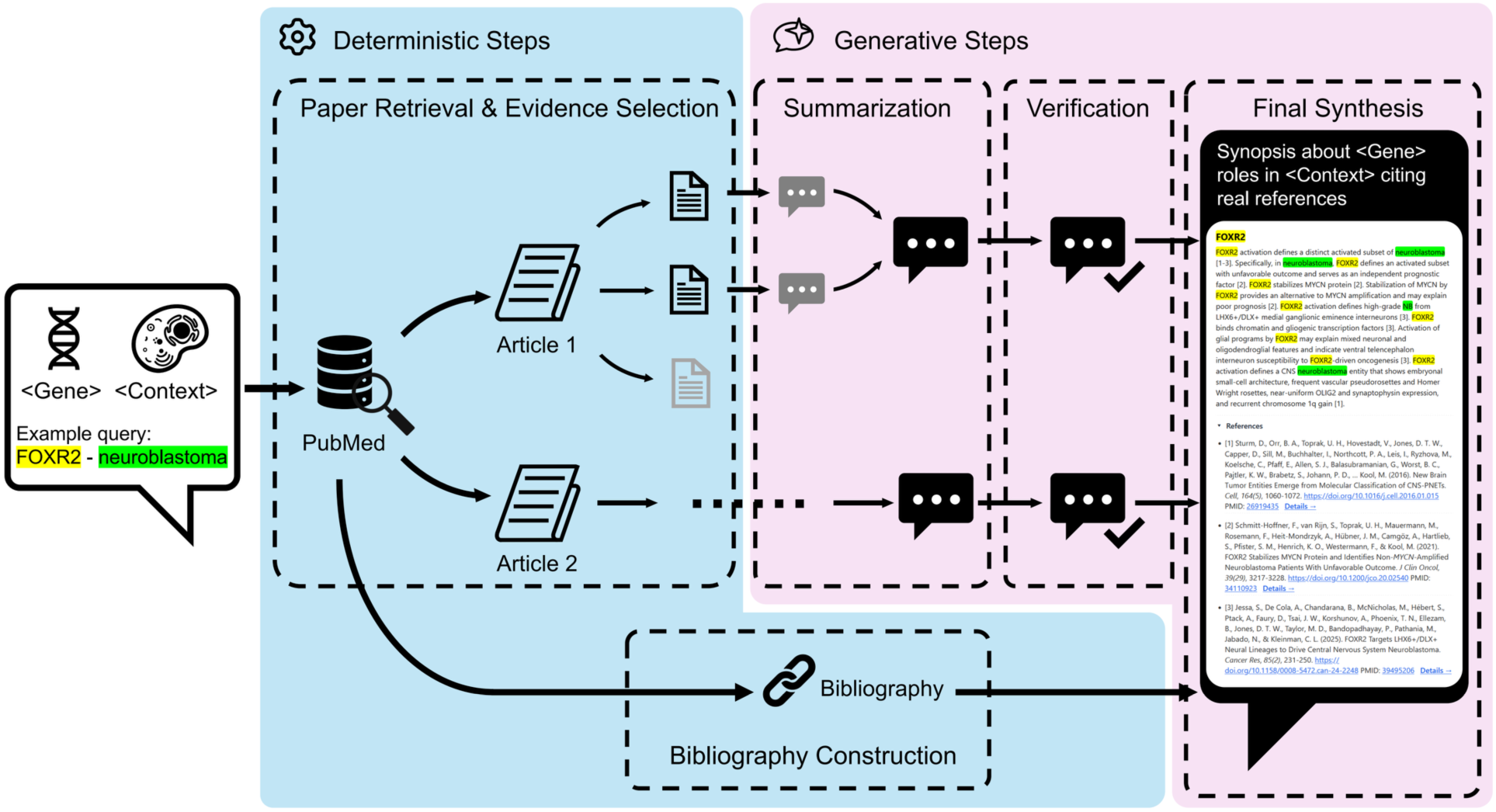
GeneKnow, an auditable AI architecture for source-grounded gene-context evidence synthesis. Schematic of the GeneKnow Discover workflow, illustrated with a representative gene-context query. Evidence-critical operations performed deterministically, including paper retrieval, passage selection, provenance tracking, and bibliography construction, are shown in blue. Generative operations constrained by retrieved evidence, including hierarchical summarization, claim-level self-verification, and cross-paper synthesis, are shown in purple. Intermediate outputs preserve traceability from the final synopsis to article-level summaries and source passages.

GeneKnow implements two complementary modes, Discover and Inspect, each initiated by a gene name and biological context. In Discover mode, GeneKnow performs a controlled academic search of the gene-context query using sources such as PubMed, identifies candidate papers, and applies deterministic procedures for evidence selection and bibliography construction. Instead of retrieving isolated text fragments from a pre-indexed vector store, GeneKnow first selects relevant papers and then ranks paragraph-level evidence in each paper by joint gene-context relevance. This paper-first strategy preserves article structure while improving exact entity grounding and reproducibility. Inspect mode begins with a user-specified paper and applies the same downstream evidence-selection and synthesis workflow.

After deterministic evidence selection, GeneKnow performs hierarchical synthesis from passage-level summaries to article-level interpretations and, in Discover mode, a final cross-paper synopsis. By concentrating each generative step on a restricted set of high-relevance evidence, this design preserves the organization of the source literature and enables compact, low-cost models with narrower context windows, such as gpt-5-mini, to perform effectively^16–18^. GeneKnow then decomposes article-level summaries into atomic claims and iteratively verifies each claim against the corresponding evidence passages, revising or removing claims that are not fully supported. Retrieved passages, verified article summaries, and final synopses are stored as separate intermediate products, preserving a traceable path from each synthesized statement to its source evidence.

### Hierarchical evidence processing and self-verification improve single-paper faithfulness

To determine which components of the framework contribute to faithfulness, we performed an ablation experiment on single-paper summarization. GeneKnow Inspect was compared with Hierarchical LLM, which retained passage selection and hierarchical summarization but omitted self-verification, and Full-paper LLM, which directly summarized the full extracted article without passage selection, hierarchical processing, or self-verification. Across 20 papers representing distinct gene-context pairs, every claim generated by GeneKnow Inspect was judged supported by the corresponding source text, whereas both ablation baselines produced evidence-uncertain or evidence-opposed claims in some cases (Fig. 2a). GeneKnow Inspect also achieved the highest alignment, coverage, and overall F1 score, followed by Hierarchical LLM and Full-paper LLM (Fig. 2b-d). These results indicate that evidence-focused hierarchical processing improves extraction from long articles and that iterative self-verification provides an additional gain in claim faithfulness.

**Figure 2.**
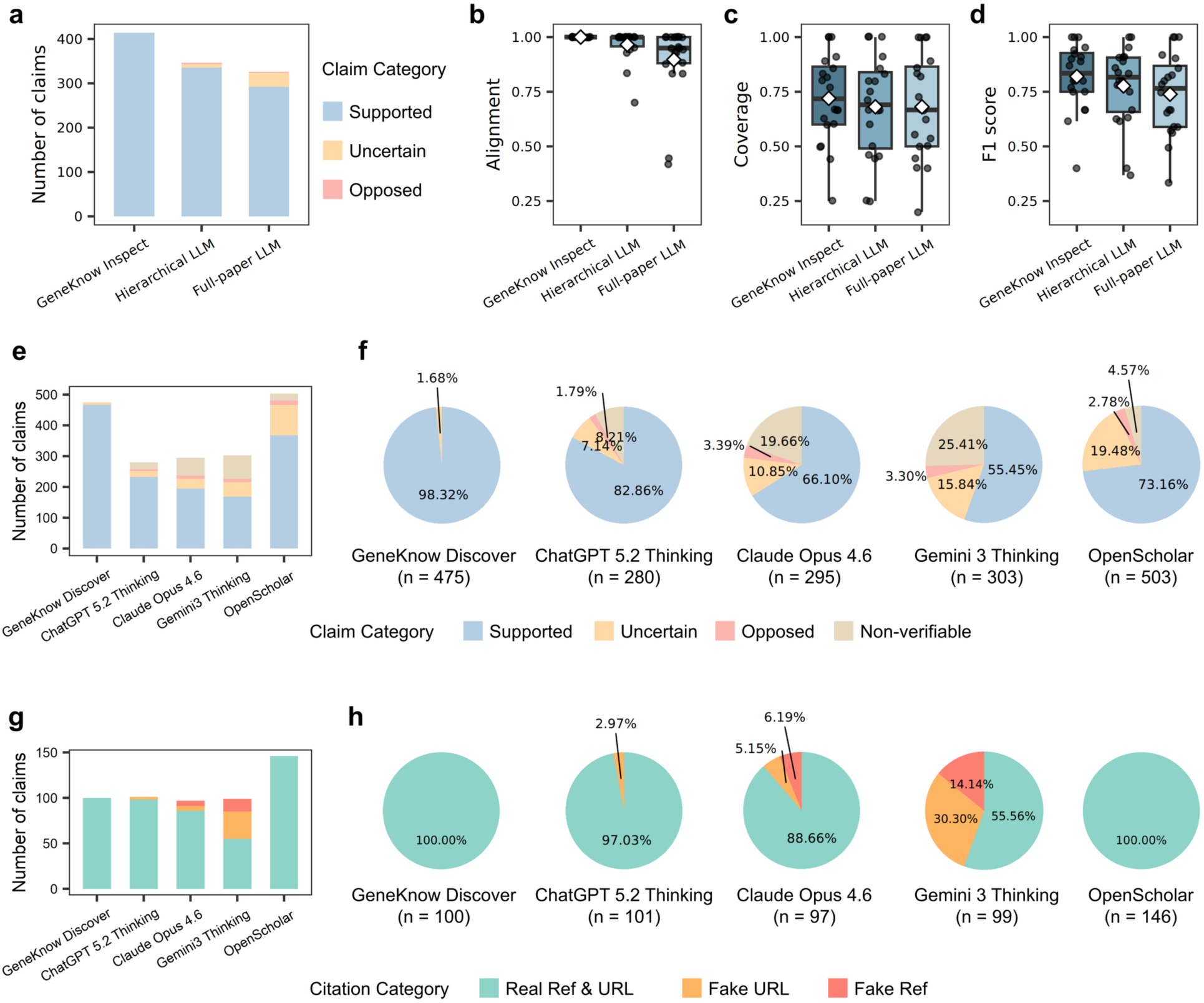
Systematic benchmarking of GeneKnow claim support, evidence coverage, and citation fidelity. a-d, Ablation analysis of GeneKnow Inspect. (a) Composition and number of claims generated by each method. Source text was examined to classify each claim as supported, opposed, or uncertain. (b-d) Alignment (b), coverage (c), and F1 score (d) across test cases. Center lines indicate medians; boxes indicate interquartile ranges (IQRs); whiskers extend to the most extreme values within 1.5 x IQR; points represent individual cases. e,f, End-to-end benchmarking of GeneKnow Discover against leading general-purpose and literature-focused AI systems. (e) Number and category composition of claims generated by each system. Claims were classified as supported, opposed, or uncertain by comparison with cited source text; claims lacking usable citations or citing fabricated references were classified as non-verifiable. (f) Proportions of claim categories for each method. g,h, Citation fidelity. (g) Number and category composition of cited references. ’Real Ref & URL’ indicates an authentic reference linked to the intended publication; ’Fake URL’ indicates an authentic reference linked to an invalid or different publication; ’Fake Ref’ indicates a fabricated reference title. (h) Proportions of citation-fidelity categories for each method.

### GeneKnow improves both claim support and evidence yield

We next tested GeneKnow Discover in an end-to-end setting that combined literature retrieval with cross-paper gene-context synthesis. We used an asking-and-answering-questions framework^19^, in which each generated claim was converted into a yes/no question and evaluated against either the claim itself or its cited source evidence. Claims were classified as supported when the evidence-derived answer matched the claim-derived answer, opposed when the source contradicted the claim, uncertain when the cited evidence was insufficient, and non-verifiable when a usable citation was absent or fabricated. We compared GeneKnow with several web-enabled general-purpose AI systems and OpenScholar^9^, a retrieval-augmented system for scientific literature question answering, across 20 prespecified gene-cell type and gene-disease relationships, such as SOX^9^ in astrocytes and FOXA^1^ in prostate cancer.

GeneKnow achieved the highest supported-claim proportion, with a large claim yield at a comparable synopsis length (Fig. 2e). Among 475 GeneKnow claims, 98.32% were supported and 1.68% were uncertain, with no opposed or non-verifiable claims. ChatGPT 5.2 Thinking, Claude Opus 4.6, and Gemini 3 Thinking achieved supported-claim proportions of 82.86%, 66.10%, and 55.45%, respectively (Fig. 2f), and each produced higher fractions of uncertain, opposed, or non-verifiable claims. Across these general-purpose systems, greater claim support generally coincided with lower claim yield, indicating a support-yield tradeoff. GeneKnow departed from this pattern by producing more claims while maintaining the highest support rate. GeneKnow also exceeded OpenScholar in claim faithfulness in this benchmark, demonstrating the value of a domain-constrained workflow for gene-context evidence synthesis.

### Deterministic bibliography construction ensures citation fidelity

GeneKnow also achieved the highest citation fidelity in the benchmark. Each extracted reference was manually examined for title authenticity and DOI or URL validity. All GeneKnow citations corresponded to authentic publications and resolved to the intended sources, yielding 100% real-reference and real-link accuracy. OpenScholar likewise achieved 100% citation authenticity, consistent with its retrieval-grounded design. By contrast, approximately 3% of links generated by ChatGPT 5.2 Thinking were incorrect, and Claude Opus 4.6 and Gemini 3 Thinking showed substantially lower real-reference and real-link accuracy (Fig. 2g,h). These findings highlight a specific advantage of GeneKnow architecture, in which deterministic bibliography construction improves citation authenticity over uncontrolled model-generating referencing.

GeneKnow has been used to curate literature-supported ground-truth evidence during development of BARTsc^20^, illustrating its practical utility for constructing traceable reference evidence in computational method development. Collectively, the ablation and end-to-end benchmarks support GeneKnow as a reproducible and source-grounded framework for gene-context evidence synthesis, with high claim support, strong evidence yield, and deterministic citation provenance in the evaluated tasks.

## Discussion

GeneKnow demonstrates a general strategy for increasing the reliability of generative AI in scientific evidence synthesis. Rather than treating an LLM as an end-to-end scientific authority, the framework assigns evidence-critical operations to deterministic procedures and restricts generative models to semantically demanding tasks whose outputs can be checked against retrieved sources. In the evaluated gene-context tasks, this division of labor supported accurate single-paper extraction, high-fidelity cross-paper synthesis, and authentic citation construction while preserving passage-level provenance. The contribution of GeneKnow is therefore not only a specialized literature tool, but also an auditable architecture for converting distributed biomedical text into inspectable biological evidence.

Several limitations define the scope of the current study. Hierarchical compression can omit source-specific details or weaken contextual qualifiers as evidence moves from passages to article summaries and final synthesis, potentially leading to over-generalization or uncertainty in the final synopsis, which is reflected by the small fraction of uncertain claims in the Discover benchmark. GeneKnow reduces this risk through iterative claim verification and a cross-stage narrative backbone centered on role in context, mechanism, and outcome, but it does not eliminate it entirely. Future improvements could further address this limitation by enhancing claim-level tracking across summarization stages. In addition, GeneKnow evaluates whether generated claims are consistent with cited sources, but it does not assess the quality and causal strength of the underlying studies. Future work could incorporate evidence grading frameworks that distinguish observational associations from causal perturbation evidence and prioritize findings supported by multiple independent studies.

GeneKnow can potentially be extended to support more complex literature queries, including functional relationships among multiple genes and nested biological contexts, such as a cell type within a disease or treatment setting. It was suggested recently that gene embeddings derived from explicit functional descriptions preserve biological structure^21^, and context-enriched gene summaries can improve tissue-specific prediction tasks^22^. Thus, another future direction is to use GeneKnow as a trustworthy data engine, and input verified GeneKnow summaries for context-conditioned gene embeddings. Overall, GeneKnow provides a practical framework for trustworthy, reproducible, and source-grounded gene-context literature synthesis for biomedical research.

## Methods

### GeneKnow framework

GeneKnow is a software framework for source-grounded gene-context literature synthesis. The system operates in two modes. Discover mode retrieves and integrates evidence across multiple papers for a specified gene-context query, whereas Inspect mode analyzes a designated paper in a single-study setting. All internal generative steps in the reported runs used gpt-5-mini through the OpenAI API with default parameters, although the framework is designed to support alternative LLM backends. All LLM-based evaluation steps used the gpt-5 model. The GeneKnow Python implementation used requests (v2.32.5) for API access, openai (v2.6.1) for model calls, pandas (v2.3.3) and numpy (v2.3.3) for tabular processing and score calculation, and jsonschema (v4.25.1) for structured-output validation. Complete prompts for passage summarization, article summarization, self-verification, brief generation, and final synopsis generation are provided in the Supplementary Material.

### Query construction, paper retrieval, and full-text inclusion

GeneKnow constructs search queries from the exact gene symbol and user-supplied context aliases. The query template joins a gene group and a context group with Boolean AND. The gene symbol is quoted exactly; optional gene aliases are joined with OR when enabled, as are optional context aliases. A typical query therefore takes the form (GENE OR GENE_ALIAS) AND (CONTEXT OR CONTEXT_ALIAS). GeneKnow can automatically expand gene aliases through NCBI Gene. PubMed is used as the default search engine, with Europe PMC and Scopus also supported.

In Discover mode, search depth is controlled by two parameters: --search-limit specifies the maximum number of candidate papers returned by the search engine, and --max-papers specifies the maximum number of PMCID-accessible full-text papers reviewed. For the reported runs, --search-limit was set to 25 and --max-papers to 5. To ensure reproducible full-text access, GeneKnow excludes candidate papers without a PMCID from Discover review. Full-text XML is downloaded from PMC when available, with Europe PMC used as a fallback when PMC retrieval fails. In Inspect mode, the target paper is specified directly by PMID or PMCID, after which the same passage-selection and synthesis workflow is applied.

### Paper preprocessing and evidence passage selection

GeneKnow parses full-text XML into paragraph-level units before LLM processing. XML parsing was implemented with beautifulsoup4 (v4.14.2) and the lxml parser (v6.0.2). Narrative sections, including the abstract, introduction, results, discussion, and conclusions, are retained for analysis, whereas figure-container text is excluded to avoid isolated panel labels or captions without sufficient narrative context. Paragraphs containing 20-500 words are retained directly; longer paragraphs are divided into sentence-based chunks that remain within the maximum length.

For each downloaded paper, GeneKnow tokenizes every valid paragraph and normalizes mentions of the queried entities. Specifically, mentions of the query gene and its aliases are replaced with the special token *t_Gene_*, and mentions of the query context and its aliases are replaced with the special token *t_Context_*. For each paragraph, GeneKnow first computes BM25 scores separately for *t_Gene_* and *t_Context_* and combines them as:

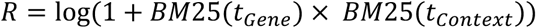

Here, *R* is the joint relevance score, which prioritizes passages containing lexical evidence for both the gene and the queried context. GeneKnow retains only passages with a positive *R* and forwards at most the top --max-passages passages from each reviewed paper to the summarization stage. Tokenization and BM25 scoring are implemented with the Python package bm25s (v0.2.14).

### Prompt-guided hierarchical summarization

GeneKnow uses a hierarchical workflow comprising passage-level summaries, article-level summaries, and, in Discover mode, a final cross-paper synopsis. Each retained passage is first summarized independently under a 100-word limit. This stage also filters passages dominated by methods, references, figure-only content, or text that does not directly address the queried gene-context relationship. The remaining passage summaries are then integrated into a single article-level summary of at most 150 words.

A central prompt-design feature is a consistent narrative backbone across stages: role in context, mechanism, and outcome. When the source evidence supports a gene-centered narrative, prompts direct the model to preserve this structure and the original relationship type, including causal, mechanistic, associative, hierarchical, or parallel relationships. When the evidence does not support such a narrative, output is restricted to direct gene-context statements without imposing a mechanistic storyline. Across all stages, prompts explicitly discourage overgeneralization, unsupported strengthening of causality, loss of experimental conditions, and restructuring of the original evidence chain.

### Iterative self-verification and revision

GeneKnow subjects each article-level summary to iterative self-verification. Atomic claims are first extracted from the draft summary and then evaluated against every retained evidence passage from the corresponding article. The verification model assigns each claim a label of ’yes,’ ’no,’ or ’partial.’ A claim is labeled ’yes’ only when the evidence preserves the stated predicate, relationship type (for example, causal, hierarchical, parallel, or associative), and relevant context (for example, species, condition, or treatment). A claim is labeled ’no’ when it conflicts with the evidence or when no relevant supporting phrase is present. Claims that are inaccurate but correctable, such as those missing an essential condition, are labeled ’partial’ and accompanied by a proposed revision. The resulting verification report records the labels and proposed revisions for each claim-evidence comparison.

GeneKnow then aggregates the verification reports into a claim-level revision plan. Claims are assigned to three mutually exclusive sets: “keep”, “deletion”, or “revision”. A claim enters the “keep” set if at least one passage returns a ’yes’ label; the “deletion” set if all passages return ’no’; and the “revision” set if at least one passage returns ’partial’ and none returns ’yes.’

For claims in the “revision” set, GeneKnow generates a corrected formulation by aggregating candidate revisions across verification reports. When multiple revisions are proposed, Jaccard similarity is used to select the candidate with the greatest lexical overlap with the original claim. This heuristic assumes that the most faithful revision is the one that preserves the highest proportion of the original terminological content while addressing the identified inaccuracy.

The revision plan is applied through a final editorial pass. Claims assigned to deletion are removed, claims assigned to revision are replaced with the selected corrected formulations, and brief transitional language is added when needed to preserve narrative coherence. Major intermediate outputs are stored separately, including evidence-passage files, article-level summaries, verification reports, final outputs, and token-usage logs.

### Final synthesis and bibliography construction

In Discover mode, each verified article-level summary is compressed into a structured brief of no more than 40 words. When supported by the evidence, the prompts retain the role in context -> mechanism -> outcome narrative backbone. Source-tagged briefs are then integrated into a unified cross-paper synopsis. The model first identifies recurrent gene roles and then presents source-specific narratives without collapsing mechanistic or relational distinctions across studies. Citations are placed at sentence endings. GeneKnow then constructs the bibliography deterministically from Europe PMC metadata, renumbers references by first citation, and appends them to the synopsis.

### Benchmark of Inspect mode

We evaluated GeneKnow Inspect using 20 prespecified gene-context cases that were also included in the Discover benchmark. The cases covered both gene-cell type and gene-disease relationships and were not used during prompt development. The complete case list and target papers are provided in the Supplementary Material. We compared three approaches: GeneKnow Inspect, which combined passage retrieval, hierarchical summarization, and self-verification; Full-paper LLM, which directly summarized extracted full-paper text without staged passage selection or verification; and Hierarchical LLM, which used the same passage selection and multi-stage summarization as GeneKnow Inspect but omitted self-verification.

An AI-driven evaluation framework was implemented following an asking-and-answering-questions approach. For alignment, atomic claims were first extracted from each system summary and converted into yes/no questions. Each question was then answered by the question-answering model using both the matched evidence text and the original claim from which the question was generated, with answer options restricted to “Yes,” “No,” and “Idk.” “Idk” was assigned only when the evaluation prompt determined that the available evidence or summary did not provide enough information to support a yes or no answer. Alignment was defined as the proportion of non-Idk claim answers for which the claim-derived answer matched the evidence-derived answer. For coverage, candidate nugget questions were generated automatically from supported claims identified in the alignment outputs. These questions were then manually curated, blinded to method identity, to remove redundancy and retain questions directly related to the gene’s major role or function in the queried context. Each summary was subsequently used to answer the shared nugget-question set, and coverage was defined as the proportion of non-Idk nugget answers recovered by the summary. The F1 score was calculated as the harmonic mean of alignment and coverage:

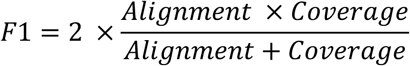

All LLM-based evaluation steps, including claim extraction, question generation, and question answering, were performed with gpt-5.

### Benchmark of Discover mode

#### Selection of comparison systems

We evaluated GeneKnow Discover on the same 20 prespecified gene-context cases used in the Inspect benchmark. The benchmark tested whether each system could generate a cross-source, evidence-grounded synopsis of the relationship between a specified gene and biological context. GeneKnow was compared with the general-purpose web-interface systems ChatGPT 5.2 Thinking, Claude Opus 4.6, and Gemini 3 Thinking, as well as OpenScholar, a retrieval-augmented system developed for scientific literature question answering.

#### Execution of web-interface LLM systems

The web-interface systems were run with browsing or web search enabled and received the same prompt requesting a concise, single-paragraph synopsis with supporting references (Supplementary Material). Each output was generated once between February and March 2026. Formatting errors and citation-generation failures were retained as observed rather than corrected by rerunning the system.

#### Execution of the OpenScholar System

OpenScholar was run using the released workflow with default hyperparameters unless otherwise specified. Each benchmark query was encoded as a JSONL record containing a query identifier, gene symbol, biological context, and raw query text. Because OpenScholar requires a short question-form input, each gene-context pair was converted into a single-sentence question asking about the functional relationship between the specified gene and context (Supplementary Material).

For retrieval, OpenScholar local search was run against the downloaded datastore-V3, and the top 100 candidate passages were retained for each query. Raw retrieval outputs were converted to the context format required by the downstream OpenScholar scripts. We then applied the released Semantic Scholar enrichment step using the OpenScholar search API utility. This step used the Semantic Scholar API and an OpenAI API call with gpt-4o to add paper-level abstracts and metadata to the locally retrieved contexts.

Final answers were generated with the released OpenScholar-8B model. Retrieved contexts were enabled, and generation used zero-shot prompting, a maximum output length of 500 tokens, no more than three passages per paper, and the top 10 contexts. We enabled the released cross-encoder reranker together with post hoc attribution, self-feedback, abstract use, normalized citations, and cross-encoder ranking. For inspection and format normalization, the resulting JSON output was converted into a readable form in which numeric citations were mapped to their corresponding passages or papers.

#### Claim-based evaluation workflow

We performed claim-based evaluation of each system output to assess both claim faithfulness and reference authenticity. Claims were extracted automatically and linked to references through in-text citations. Each claim was converted into a yes/no question, and an evaluator model answered the question using either the generated claim or the cited source evidence. A claim was classified as supported when the evidence-derived answer matched the claim-derived answer, opposed when the source evidence contradicted the claim, and uncertain when the cited evidence was insufficient to determine correctness. Claims without a usable citation or with a fabricated reference were classified as non-verifiable.

Reference authenticity and DOI or URL validity were reviewed manually. A reference was considered authentic when its cited title matched a real publication. A link was considered valid when the DOI or URL resolved to the same publication. When a cited paper was authentic but the original link was invalid or inaccessible, an alternative valid link was recorded for downstream evidence verification.

Cited papers were manually collected as HTML files or as Markdown files converted from PDF with PyMuPDF4LLM (v0.3.4). The same section-filtering procedure used by GeneKnow was applied to extract the main text. Each claim was then verified with an asking-and-answering-questions framework analogous to that used in the Inspect benchmark, except that the cited paper or papers served as the evidence. A claim was classified as supported if at least one authentic and accessible cited paper verified it; opposed if the evidence-derived answer contradicted the claim-derived answer; uncertain if the evidence-derived answer was ’Idk’; and non-verifiable if the claim lacked a usable citation or cited a fabricated reference. Visualizations were generated in R with ggplot2 (v4.0.2), dplyr (v1.2.0), tidyr (v1.3.2), and cowplot (v1.2.0).

## Code Availability

The GeneKnow source code is freely available at https://github.com/zang-lab/GeneKnow

## Acknowledgments

H.Z. is a Wagner Fellow at the University of Virginia (UVA) School of Medicine and receives a Farrow Fellowship from the UVA Comprehensive Cancer Center. This work was supported in part by US National Institutes of Health grant R35GM133712 (C.Z.).

## Notes

https://github.com/zang-lab/GeneKnow

### Competing Interest Statement

The authors have declared no competing interest.

### Summary of Updates

Figure 2 have been revised by adding OpenScholar in the benchmarking. Manuscript has been revised with in-depth discussion and improved clarity. Supplementary materials have been updated with more information for enhanced transparency and reproducibility.

