## Supplementary Material for "GeneKnow: An auditable AI framework for source-grounded biological evidence synthesis"

### GeneKnow: AI-powered literature synthesis for gene-context analysis

This document catalogues all structured output schemas, LLM prompts and test cases used in the development of GeneKnow.

### 1. Output Schemas

#### 1.1. CLAIM_SCHEMA

Defines an array of atomic claims with sequential integer IDs and claim text strings.

```
{
 "title": "ClaimsObject",
 "type": "array",
 "items": {
   "type": "object",
   "required": ["id", "claim"],
   "additionalProperties": false,
   "properties": {
     "id": {"type": "integer"},
     "claim": {"type": "string"}
   }
 }
}
```

#### 1.2. VERI_SCHEMA

Defines a verification report array with support flags (yes/no/partial) and optional revised claims.

```
{
 "title": "VeriObject",
 "type": "array",
 "items": {
   "type": "object",
   "required": ["id", "flag", "new"],
   "additionalProperties": false,
   "properties": {
     "id": {"type": "integer"},
     "flag": {"type": "string", "enum": ["yes", "no", "partial"]},
     "new": {"type": "string"}
   }
 }
}
```

#### 1.3. CATEGORY_SCHEMA

Defines an array of categorized claims assigning each to one of five biomedical categories.

```
{
  "title": "CategoryObject",
  "type": "array",
  "items": {
    "type": "object",
    "required": ["id", "claim", "category"],
    "additionalProperties": false,
    "properties": {
      "id": {"type": "integer"},
      "claim": {"type": "string"},
      "category": {"type": "string", "enum": ["role", "context", "mechanism", "outcome", "other"]}
    }
  }
}
```

#### 1.4. BRIEF_SCHEMA

Defines a brief summary object with a structural type (`logical` or `factual`) and the summary text.

```
{
  "title": "BriefObject",
  "type": "object",
  "required": ["type", "brief"],
  "additionalProperties": false,
  "properties": {
    "type": {"type": "string", "enum": ["logical", "factual"]},
    "brief": {"type": "string"}
  }
}
```

### 2. Function Prompts

#### 2.1. summarize_passage

Summarizes the gene–context relationship from a single passage.

##### System Prompt

You are a concise academic assistant. You summarize ONLY from the provided context. Do not add facts or world knowledge. Use plain English, avoid speculation.

##### Developer Prompt

Instructions:
- Write a ≤100-word summary of the relationship between {gene} and {context} using only the provided text.
- Prioritize high-level statements and the most specific role first (e.g., specifies, required for,

drives identity).
- First determine the relationship structure in the passage (causal chain, intermediary mechanism, parallel listing, or association), then restate it faithfully without merging steps, strengthening claims, or changing hierarchy. Choose connectors based on structure (not fluency); if unsure, use neutral wording.
- Preserve key context and conditions explicitly stated (e.g., developmental stage, disease/injury state, species, tissue/region, experimental setting). Do not add, assume, or generalize beyond the text.
Content Rules:
- First determine whether the passage is {gene}-centered (systematically elaborates {gene} function in {context}) or non–{gene}-centered (mentions {gene} only peripherally or in independent facts).
• If non–{gene}-centered: summarize only direct statements that explicitly mentions both {gene} and {context}, excluding indirect, background, or contextual information.
• If {gene}-centered: write a summary following a logical flow: role(context) → mechanism → outcome, preserving exact relationship structure and drivers without merging steps.
- Inforamtion to ignore: facts not directly related to {gene}, Assay/protocol details, long gene lists, Over-precise numbers
Output:
- Write in natural prose, NO title, NO subtitles, NO bullet points, No arrows.
- AVOID comma splice, MAX two clear clauses per sentence.
- If the passage is methods/figure legend/references, or no {gene}–{context} relationship is present, output: no function revealed

**User Prompt**

Read the following passage and summarize the relationship between {gene} and {context}.

{passage}

#### 2.2. summarize_article

Merges multiple passage summaries into one article-level summary.

**System Prompt**

You are a concise academic assistant. You summarize ONLY from the provided context. Do not add facts or world knowledge. Use plain English, avoid speculation.

**Developer Prompt**

Instructions:
- Merge summaries split by '-' into a single-paragraph summary, remove redundancy (<150 words).
- Focus on claims involving {gene} and {context},
- Stay faithful to the text: do not add or infer details; keep explicit conditions and contexts when stated (e.g. “stage”, “disease”, “model”). You may reorganize for clarity.
Structure:
- First determine whether any passage is a systematic elaboration of {gene} role (follows a logical flow of role->mechanism->outcome), or all the passages just state {gene}-related facts.

• If systematic: write a summary following a logical flow: role(context) → mechanism → outcome, preserving exact relationship structure and drivers without merging steps.

Content Rules:
- Prioritize Core Biological Claims about {gene} over secondary details.
- Do not merge findings across distinct contexts unless explicitly presented together.
- Inforamtion to ignore: facts not directly related to {gene}, Assay/protocol details, long gene lists, Over-precise numbers
Writing Constraints:
- Use causal verbs only if explicitly supported by the text.
- AVOID comma splice, MAX two clear clauses per sentence.
- Use explicit subjects and avoid ambiguous pronouns or demonstratives such as “these cells", "this mechanism".
- No title, No subtitles, Return plain text only.
- If the passages only non–{gene}-centered content, output: no function revealed

**User Prompt**

Merge the following summaries together into a single summary

{passage_summaries_joined_with_newlines}

#### 2.3. extract_claims

Decomposes a summary text into atomic claims for verification.

**System Prompt**

You are a claim extractor. Extract atomic, checkable claims that (1) preserve the original meaning, (2) add no external info, (3) avoid redundancy/overlap, and (4) collectively cover the entire source.

**Developer Prompt**

A claim is defined as one proposition anchored to a single subject–predicate relationship.
Instructions:
- Extract atomic, checkable claims that exactly reflect the source—no external facts, inference, or speculation.
- Preserve hedging/negation/conditions, entity names/aliases as written, and any numbers, units, directions, or temporal cues.
- Replace all pronouns and demonstratives such as "this population", "these cells" with the full explicit entity name described in the source text.
- Split long or compound sentences into multiple claims.
- Each claim must be self-contained and interpretable in isolation, restate necessary constraints, context, and entity names explicitly
- Use the fewest non-overlapping claims needed to fully cover the text.
- Return the extracted claims with sequential ids in a JSON array, example:
```
[
  {
    "id" : 1,
```

```
        "claim" : "claim1"
    },
    {
        "id" : 2,
        "claim" : "claim2"
    },
    {
        "id" : 3,
        "claim" : "claim3"
    }
]
```
- Retrun the following blank JSON array, if no claims was extracted or the source text is blank:
[]

**User Prompt**

Extract checkable claims from the following summary

Summary to extract: {summary}

#### 2.4. verify_claims_against_passage

Checks claims for semantic faithfulness to a passage.

**System Prompt**

You are acdemic fact checker. You verify the authenticity and accuracy of claims based on the given evidence. Do not add facts or world knowledge that are not mentioned by the evidence

**Developer Prompt**

Instructions:
- Read the evidence passage between <evidence> and </evidence>
- Read a JSON format claim list between <JSON> and </JSON>
- Evaluate each claim for strict semantic faithfulness to the evidence.

Judging Rules (STRICT):
- A claim is "yes" (supported) ONLY if:
1. All relationships (causal, hierarchical, parallel, associative) match the evidence exactly.
2. The strength of verbs and modality (e.g., involved vs mediates) is preserved.
3. No extra conditions, scope, timing, or context are added.
4. No restructuring of relationships occurs.
- A claim is "partial" if ANY of the following occur:
1. Overstatement: The claim strengthens the evidence wording.
2. Relationship Distortion: The claim changes structure:
    parallel → hierarchical
    listing → causal
    associative → mechanistic
3. Scope/Context Drift: added extra contexts, or lacks necessary contexts
4. Minor Inaccuracy: mostly correct but slightly wrong wording or ambiguous subject reference.
- A claim is "no" if:

1. meaning conflicts with evidence
2. the evidence does not contain any relevant phrases of the claim
- If one claim is unsupported, or completely supported without any necessay revisions, do not return a new claim.
- Avoid making inferences based on anything other than the evidence.
- Return the claim ids, support judgement, and revised claims in a JSON array,
template:
[{"id": <same>, "flag": <yes or no or partial>, "new": <new claim>},...]
example:

```
[
{
   "id" : 1,
   "flag" : "yes",
   "new": ""
},
{
   "id" : 2,
   "flag" : "no",
   "new": ""
},
{
   "id" : 3,
   "flag" : "partial",
   "new": "Gene A is specifically expressed in cell type B during early development"
}
]
```

- Retrun the following blank JSON array, if no claims was extracted or the source text is blank:
[]

**User Prompt**

Identify whether the claims are completely/partially/not supported by the evidence

<JSON>
{claims_json}
</JSON>

<evidence>
{passage}
</evidence>

#### 2.5. apply_revisions_to_summary

Edits a summary by applying deletions and replacements.

**System Prompt**

You are a concise academic assistant. You edit a given text based on instructions. Use plain English, avoid speculation. Do not add facts or world knowledge.

**Developer Prompt**

Instructions:
- Read A TEXT block between <text> and </text>, this is the document to edit.
- Read A JSON array of revisions between <revisions> and </revisions>: [{"old": "...", "new": ""} or {"old": "...", "new": "..."}, ...]
- For entries with empty "new": DELETE the old claim from the text
- For entries with non-empty "new": REPLACE the old claim with the new claim
- Maintain the fluency and clarity in the writing, bridge gaps with brief transitions if necessary.
- Only return edited text in plain text. Do not add any comments or section titles.
- If no revisions are provided, maintain the original text.

**User Prompt**

Apply the specified revisions to the given text.

<text>
{summary}
</text>

<revisions>
{revision_list_json}
</revisions>

#### 2.6. categorize_claims

Classifies claims into role, context, mechanism, outcome, or other.

**System Prompt**

You are a biomedical claim categorizer. You analyze scientific claims and classify them into predefined categories based on their content. Be precise and consistent in your categorizations.

**Developer Prompt**

Instructions:
- Read the JSON format claim list provided by the user.
- Categorize each claim into ONE of the following categories:
  1. "role" - The biological role, function, or identity of the gene (e.g., "Gene X acts as a transcription factor", "Gene X is a key regulator of Z", "Gene W functions as a kinase")
  2. "context" - Experimental or biological context, conditions, or settings, including context-specific role (e.g., "during embryonic development", "in the adult hippocampus", "under hypoxic conditions", "in cancer cells")
  3. "mechanism" - Molecular or cellular mechanisms, interactions, or processes (e.g., "activates Wnt signaling pathway", "interacts with Beta-catenin", "regulates gene expression through chromatin remodeling", "phosphorylates target protein")
  4. "outcome" - Downstream effects, phenotypic consequences, or other observations (e.g., "promotes neuronal differentiation", "leads to apoptosis", "results in increased cell proliferation")
  5. "other" - Claims that don't fit any of the above categories or are primarily methodological, background information, etc.

- Preserve the original claim text and IDs exactly.
- Return the results as a JSON array with the following format:

```
[
  {
    "id": 1,
    "claim": "original claim text",
    "category": "role"
  },
  {
    "id": 2,
    "claim": "original claim text",
    "category": "mechanism"
  }
]
```

- Return an empty JSON array [] if no claims are provided or the input is blank.
- Ensure each claim is assigned exactly one category.

**User Prompt**

Categorize the following claims:

{final_claims_json}

#### 2.7. generate_brief_from_summary

Generates a ≤40-word brief summary from an article summary.

**System Prompt**

You are a biomedical synthesis writer. You create concise, flowing summaries from scientific texts. Focus on capturing the essential narrative in minimal words.

**Developer Prompt**

Instructions:
- Read the provided article summary.
- Determine structure type: whether the summary follows a {gene}-centered logical flow of role(context) → mechanism → outcome (logical), or only present discrete facts without logical connections (factual).
- Rephrase into one paragraph of clear brief summary under 40 words based on the structure type:
    - If logical type: preserve the role(context) → mechanism → outcome writing structure.
    - If factual type: summarize only direct statements that explicitly mention both {gene} and {context}, excluding indirect, background, or contextual information.

Content Rules:
- Avoid over-compression, focus on the cole biological claims about {gene}
- Include at most 1-2 directly involved mechnasims
- Exclude secondary mechanisms, secondary outcomes, Assay/protocol details, long gene lists, Over-precise numbers
- Claims across distinct contexts or conditions must be expressed as separate statements.

Faithfulness to original relationship
- First determine the relationship structure expressed in the passage (causal chain, intermediary mechanism, parallel listing, or association)
- Restate the structure faithfully without collapsing steps, changing hierarchy or adding links
- choose connectors strictly based on structure, not fluency.
- Ensure predicates expressing relationships remain unchanged (e.g. specify, regulate, upregulate, mediate, coactivate)
Causality control
- Do not introduce causal relationship unless they are directly supported by the input.
- Use causality-implying words cautiously, including (e.g. "underlie", "drive", "induce", "to", "through", "by")
Writing Constraints:
- Write in natural prose, NO title, NO subtitles, NO bullet points, No arrows.
- AVOID comma splice, MAX two clear clauses per sentence.
- Ensure every sentence is self-contained and unambiguous, without parentheses or supplemental information.
- Use explicit subjects and avoid ambiguous pronouns or demonstratives such as "these cells", "this mechanism".
Output Format:
Return a JSON object with two keys:
- "type": "logical"/"factual"
- "brief": the brief summary text
Example:
{"type": "logical", "brief": "Gene X acts as a transcription factor that activates pathway Y, leading to cell differentiation."}

**User Prompt**

Generate a brief summary from the following article summary:

{summary}

#### 2.8. generate_synopsis_from_briefs

Synthesizes briefs into a synopsis with in-text citations.

**System Prompt**

You are a biomedical synthesis writer. You synthesize multiple brief summaries about {gene} in {context} into a coherent synopsis paragraph with proper citations. Use the original IDs provided for citations.

**Developer Prompt**

Definition of core role:
- The most general, context-robust function {gene} plays — the high-level biological job it performs across multiple studies, especially in the context of {context}.
Input Format:
- Each brief has: "id" (for citations), "type" ("logical" or "factual"), and "brief" (the summary text)
Instructions:

- Read the provided JSON array of brief summaries about the relationship between {gene} and {context}. Brief summaries that do not mention {gene}-{context} relationship should be skipped.
Step1: State Core Roles
    - Identify core roles of {gene} appearing in two or more briefs.
    - Exclude mechanisms, conditions, or outcomes in this section.
    - Do NOT merge distinct roles into a single statement.
    - Skip if no core role was found
Step2: Present Context-specific Narratives
    - Prioritize "logical" type briefs and include "factual" briefs after them,
    - Present brief-specific narratives or facts about {gene}, one at a time without merging across briefs.
    - Do not skip or simplify any logical steps in the original briefs, unless already mentioned as core roles.
    - Preserve relationship types exactly (causal, mechanistic, parallel, associative), Do NOT strengthen verbs or restructure relationships.
    - Do NOT combine mechanisms or outcomes across different briefs unless explicitly shared.
Step3: Add minimal transition words to improve flow, but do NOT change meaning, relationships, emphasis, or introduce new information.
Citation Rules:
- Use in-text citations at sentence end (before period): single [1]; multiple [1][2][3].
- Use the EXACT "id" from the input JSON.
- If sources disagree, present conflicting claims in separate sentences.
Writing Constraints:
- AVOID comma splice, MAX two clear clauses per sentence.
- Use explicit subjects and avoid ambiguous pronouns or demonstratives such as "these cells", "this mechanism".
- Write a paragraph in natural prose, NO title, NO subtitles, NO bullet points, No arrows.
- Output plain text only.
- If NO valid briefs provided, return an empty string.

**User Prompt**

Synthesize the following briefs about {gene} in {context} into a synopsis with citations:

{briefs_json}

### 3. Web-interface Prompts

Write a concise single-paragraph synopsis (~200 words) describing the functional relationship between {gene} and {context}, including only claims directly supported by peer-reviewed research articles; do not infer, speculate, or generalize beyond what is explicitly reported. For each claim, cite a real publication that directly supports that specific statement, and present the references after the paragraph in APA 7th format with DOI links. If no direct evidence exists, explicitly state that instead of extrapolating.

### 4. OpenScholar Query

What functional relationships between {gene} and {cell type} have been directly reported in peer-reviewed research articles?

### 5. Test Cases

The same 20 gene-context pairs are used for both the benchmarks of GeneKnow Inspect and GeneKnow Discover. The PMID column lists the designated papers used for the benchmark of GeneKnow Inspect.

| id | gene | context | PMID |
|---|---|---|---|
| 1 | SOX9 | astrocyte astrocytic | 41271638 |
| 2 | PU.1 | macrophage | 40673490 |
| 3 | PAX5 | B cell | 39424985 |
| 4 | FOXP3 | Treg “regulatory T cell” | 36569832 |
| 5 | MYOD1 | “muscle cell” “muscle fiber” myocyte | 36739949 |
| 6 | NEUROD1 | neuron “nerve cell” | 39105169 |
| 7 | NKX2-1 | “AT2 cell” “alveolar type II cell” “alveolar type 2 cell” | 39284798 |
| 8 | RUNX2 | osteoblast | 39337587 |
| 9 | OLIG2 | oligodendrocyte oligodendroglia OPC oligodendro | 34172044 |
| 10 | CEBPA | neutrophil granulocyte | 24584857 |
| 11 | NOTCH1 | leukemia AML ALL CML CLL | 40009499 |
| 12 | PD-1 | melanoma | 30089911 |
| 13 | PD-L1 | “non-small cell lung cancer” NSCLC | 35159131 |
| 14 | KLF4 | “colorectal cancer” CRC | 37031197 |
| 15 | FOXA1 | “prostate cancer” | 32946061 |
| 16 | ESR1 | “breast cancer” | 32289273 |
| 17 | APOE4 | “Alzheimer's disease” “Alzheimer disease” AD | 29861287 |
| 18 | TCF7L2 | “diabetes mellitus” diabetes T2D T2DM | 34016596 |
| 19 | FOXR2 | neuroblastoma NB-FOXR2 | 26919435 |
| 20 | PAX3-FOXO1 | rhabdomyosarcoma | 37968277 |

### 6. Input and Output Example

#### 6.1. Input Example

Gene: FOXR2
Context: neuroblastoma NB-FOXR2

#### 6.2. Output Example

FOXR2 functions as an oncogenic driver that is essential in neuroblastoma [1,2]. FOXR2 activation designates a distinct CNS neuroblastoma entity (NB-FOXR2) [1,3,4]. FOXR2 activation designates a CNS neuroblastoma entity with embryonal small-cell histology and near-uniform OLIG2 and synaptophysin expression [1]. Sequencing identified complex genomic rearrangements converging on FOXR2 that increased its expression [1]. FOXR2 activation instigates oncogenic activity [1]. FOXR2 stabilizes MYCN protein and thereby forms an alternative mechanism to MYCN amplification [2]. The FOXR2-activated subset shows an unfavorable clinical outcome [2]. FOXR2 activation defines NB-FOXR2 as tumors from

LHX6+/DLX+ medial ganglionic eminence interneuron lineages [5]. FOXR2 binds chromatin and transcription factors to activate glial programs that may explain the tumors' mixed neuronal and oligodendroglial features [5]. FOXR2 activation is regarded as pathognomonic for CNS neuroblastoma (NB-FOXR2) [3]. All CNS NB-FOXR2 were cerebral cortical tumors [3]. CNS Neuroblastoma with FOXR2 activation (CNS NB-FOXR2) is a distinct tumor group identified among previously diagnosed CNS-PNETs [4].

**References**

[1] Sturm, D., Orr, B. A., Toprak, U. H., Hovestadt, V., Jones, D. T. W., Capper, D., Sill, M., Buchhalter, I., Northcott, P. A., Leis, I., Ryzhova, M., Koelsche, C., Pfaff, E., Allen, S. J., Balasubramanian, G., Worst, B. C., Pajtler, K. W., Brabetz, S., Johann, P. D., Sahm, F., Reimand, J., Mackay, A., Carvalho, D. M., Remke, M., Phillips, J. J., Perry, A., Cowdrey, C., Drissi, R., Fouladi, M., Giangaspero, F., Łastowska, M., Grajkowska, W., Scheurlen, W., Pietsch, T., Hagel, C., Gojo, J., Lötsch, D., Berger, W., Slavc, I., Haberler, C., Jouvet, A., Holm, S., Hofer, S., Prinz, M., Keohane, C., Fried, I., Mawrin, C., Scheie, D., Mobley, B. C., Schniederjan, M. J., Santi, M., Buccoliero, A. M., Dahiya, S., Kramm, C. M., von Bueren, A. O., von Hoff, K., Rutkowski, S., Herold-Mende, C., Frühwald, M. C., Milde, T., Hasselblatt, M., Wesseling, P., Rößler, J., Schüller, U., Ebinger, M., Schittenhelm, J., Frank, S., Grobholz, R., Vajtai, I., Hans, V., Schneppenheim, R., Zitterbart, K., Collins, V. P., Aronica, E., Varlet, P., Puget, S., Dufour, C., Grill, J., Figarella-Branger, D., Wolter, M., Schuhmann, M. U., Shalaby, T., Grotzer, M., van Meter, T., Monoranu, C. M., Felsberg, J., Reifenberger, G., Snuderl, M., Forrester, L. A., Koster, J., Versteeg, R., Volckmann, R., van Sluis, P., Wolf, S., Mikkelsen, T., Gajjar, A., Aldape, K., Moore, A. S., Taylor, M. D., Jones, C., Jabado, N., Karajannis, M. A., Eils, R., Schlesner, M., Lichter, P., von Deimling, A., Pfister, S. M., Ellison, D. W., Korshunov, A., & Kool, M. (2016). New Brain Tumor Entities Emerge from Molecular Classification of CNS-PNETs. *Cell*, 164(5), 1060-1072. https://doi.org/10.1016/j.cell.2016.01.015 PMID: 26919435

[2] Schmitt-Hoffner, F., van Rijn, S., Toprak, U. H., Mauermann, M., Rosemann, F., Heit-Mondrzyk, A., Hübner, J. M., Camgöz, A., Hartlieb, S., Pfister, S. M., Henrich, K. O., Westermann, F., & Kool, M. (2021). FOXR2 Stabilizes MYCN Protein and Identifies Non-*MYCN*-Amplified Neuroblastoma Patients With Unfavorable Outcome. *J Clin Oncol*, 39(29), 3217-3228. https://doi.org/10.1200/jco.20.02540 PMID: 34110923

[3] Siskar, A. N., Hanzlik, E., Cardenas, M. F., Eldomery, M. K., Pinto, S., Tinkle, C. L., Zhang, Q., Li, X., Lin, T., Dhanda, S. K., Reis, G., Li, D., Raghavan, R., Vortmeyer, A., Karajannis, M. A., Robinson, G. W., Onar-Thomas, A., Blackburn, P. R., Wheeler, D. A., & Chiang, J. (2025). FOXR2 activation is not exclusive of CNS neuroblastoma. *Neuro Oncol*, 27(7), 1801-1812. https://doi.org/10.1093/neuonc/noaf076 PMID: 40237561

[4] Łastowska, M., Trubicka, J., Sobocińska, A., Wojtas, B., Niemira, M., Szałkowska, A., Krętowski, A., Karkucińska-Więckowska, A., Kaleta, M., Ejmont, M., Perek-Polnik, M., Dembowska-Bagińska, B., Grajkowska, W., & Matyja, E. (2020). Molecular identification of CNS NB-FOXR2, CNS EFT-CIC, CNS HGNET-MN1 and CNS HGNET-BCOR pediatric brain tumors using tumor-specific signature genes. *Acta Neuropathol Commun*, 8(1), 105. https://doi.org/10.1186/s40478-020-00984-9 PMID: 32650833

[5] Jessa, S., De Cola, A., Chandarana, B., McNicholas, M., Hébert, S., Ptack, A., Faury, D., Tsai, J. W., Korshunov, A., Phoenix, T. N., Ellezam, B., Jones, D. T. W., Taylor, M. D., Bandopadhayay, P., Pathania, M., Jabado, N., & Kleinman, C. L. (2025). FOXR2 Targets LHX6+/DLX+ Neural Lineages to Drive Central Nervous System Neuroblastoma. *Cancer Res*, 85(2), 231-250. https://doi.org/10.1158/0008-5472.can-24-2248 PMID: 39495206

### 7. Example of claim-based evaluation

#### 7.1. Synopsis to evaluate

The following synopsis and reference list about FOXR2 function in neuroblastomas were generated by Gemini 3 thinking.

FOXR2 acts as a potent oncogenic driver in a subset of neuroblastomas, where its expression is typically mutually exclusive with $MYCN$ amplification (Schmitt-Hoffner et al., 2021). Functionally, the FOXR2 protein binds to and stabilizes the MYCN protein, protecting it from proteasomal degradation and thereby facilitating its transcriptional activity in tumors without $MYCN$ gene amplification (Schmitt-Hoffner et al., 2021). In sympathetic neuroblastoma, FOXR2 expression is significantly enriched in advanced clinical stages and identifies a subset of patients with unfavorable outcomes (Schmitt-Hoffner et al., 2021). Beyond sympathetic tumors, FOXR2 activation via enhancer hijacking is a defining genetic hallmark of central nervous system (CNS) neuroblastoma, a distinct molecular entity (Liu et al., 2022; Schmitt-Hoffner et al., 2021). In these CNS tumors, FOXR2 specifically targets $LHX6+/DLX+$ neural lineages derived from the medial ganglionic eminence, where it activates ETS transcriptional networks to drive oncogenesis (Jessa et al., 2024). This activation promotes cellular proliferation and influences mitogen-activated protein kinase (MAPK) signaling (Yang et al., 2025). Furthermore, FOXR2 directly binds to key transcription factors that coordinate the initiation of gliogenesis, suggesting a role in disrupting normal neural differentiation during tumor development (Jessa et al., 2024).

### 7.2. Table of Extracted References

| In-Text Citation | Reference | URL | Real Reference | Real URL | Alternative URL | Hash ID |
|---|---|---|---|---|---|---|
| (Schmitt-Hoffner et al., 2021) | Schmitt-Hoffner, F., van Rijn, S., Toprak, U. H., Mauermann, M., Rosemann, F., Heit-Mondrzyk, A., Hübner, J.-M., Camgöz, A., Hartlieb, S., Pfister, S. M., Henrich, K.-O., Westermann, F., & Kool, M. (2021). FOXR2 stabilizes MYCN protein and identifies non–MYCN-amplified neuroblastoma patients with unfavorable outcome. *Journal of Clinical Oncology*, *39*(29), 3217–3228. [https://doi.org/10.1200/jco.20.02540](https://www.google.com/search?q=https://doi.org/10.1200/jco.20.02540) Cited by: 41 | https://doi.org/10.1200/jco.20.02540 | TRUE | TRUE | | GSRzoKNUA_szgfUv |
| (Liu et al., 2022) | Liu, A. P. Y., Dhanda, S. K., Lin, T., Sioson, E., Vasilyeva, A., Gudenas, B., Tatevossian, R. G., Jia, S., Neale, G., Bowers, D. C., Hassall, T., Partap, S., Crawford, J. R., Chintagumpala, M., Bouffet, E., McCowage, G., Broniscer, A., Qaddoumi, I., Armstrong, G., ... Chiang, J. (2022). Molecular classification and outcome of children with rare CNS embryonal tumors: results from St. Jude Children's Research Hospital including the multi-center SJYC07 and SJMB03 clinical trials. *Acta Neuropathologica*, *144*(4), 733–746. [https://doi.org/10.1007/s00401-022-02484-7](https://doi.org/10.1007/s00401-022-02484-7) Cited by: 28 | https://doi.org/10.1007/s00401-022-02484-7 | TRUE | TRUE | | yg2ktk7SgiICsa-g |
| (Jessa et al., 2024) | Jessa, S., De Cola, A., Chandarana, B., McNicholas, M., Hébert, S., Ptack, A., Faury, D., Tsai, J. W., Korshunov, A., Phoenix, T. N., Ellezam, B., Jones, D. T. W., Taylor, M. D., Bandopadhayay, P., Pathania, M., Jabado, N., & Kleinman, C. L. (2024). FOXR2 targets LHX6+/DLX+ neural lineages to drive central nervous system neuroblastoma. *Cancer Research*, *85*(2), 231–250. [https://doi.org/10.1158/0008-5472.can-24-2248](https://doi.org/10.1158/0008-5472.can-24-2248) Cited by: 2 | https://doi.org/10.1158/0008-5472.can-24-2248 | TRUE | TRUE | | paj5mxPEndxt-zp4 |

| In-Text Citation | Reference | URL | Real Reference | Real URL | Alternative URL | Hash ID |
|---|---|---|---|---|---|---|
| (Yang et al., 2025) | Yang, P., Mahaki, H., Kalhor, N., Dirbaziyan, A., Al-Musawi, S., Sheykhhasan, M., Heidari, R., Chamanara, M., Dama, P., Ahmadieh-Yazdi, A., Manoochehri, H., & Tanzadehpanah, H. (2025). FOXR2 in cancer development: emerging player and therapeutic opportunities. *Oncology Research*, *33*, 283–300. [https://doi.org/10.32604/or.2024.052939](https://www.google.com/search?q=https://doi.org/10.32604/or.2024.052939)<br>Cited by: 7 | https://doi.org/10.32604/or.2024.052939 | TRUE | TRUE | | vRrDBwJ74nwj7Yab |

### 7.3. Table of Extracted Claims and Claim-Based Evaluation Results

| ID | Claim | Citation | Question | Claim Answer | Paper Answer | Answer Note |
|---|---|---|---|---|---|---|
| 1 | FOXR2 acts as a potent oncogenic driver in a subset of neuroblastomas. | (Schmitt-Hoffner et al., 2021) | Does FOXR2 act as a potent oncogenic driver in a subset of neuroblastomas? | Yes | Yes | |
| 2 | In a subset of neuroblastomas, FOXR2 expression is typically mutually exclusive with MYCN amplification. | (Schmitt-Hoffner et al., 2021) | Is FOXR2 expression typically mutually exclusive with MYCN amplification in a subset of neuroblastomas? | Yes | Yes | |
| 3 | The FOXR2 protein binds to the MYCN protein. | (Schmitt-Hoffner et al., 2021) | Does the FOXR2 protein bind the MYCN protein? | Yes | Yes | |
| 4 | The FOXR2 protein stabilizes the MYCN protein. | (Schmitt-Hoffner et al., 2021) | Does FOXR2 stabilize the MYCN protein? | Yes | Yes | |
| 5 | The FOXR2 protein protects the MYCN protein from proteasomal degradation. | (Schmitt-Hoffner et al., 2021) | Does FOXR2 protect MYCN from proteasomal degradation? | Yes | Idk | Passage doesn't mention proteasomal degradation specifically |
| 6 | FOXR2 facilitates MYCN transcriptional activity in tumors without MYCN gene amplification. | (Schmitt-Hoffner et al., 2021) | Does FOXR2 facilitate MYCN transcriptional activity in tumors without MYCN amplification? | Yes | Yes | |
| 7 | In sympathetic neuroblastoma, FOXR2 expression is significantly enriched in advanced clinical stages. | (Schmitt-Hoffner et al., 2021) | Is FOXR2 expression enriched in advanced stages of sympathetic neuroblastoma? | Yes | Yes | |
| 8 | In sympathetic neuroblastoma, FOXR2 expression identifies a subset of patients with unfavorable outcomes. | (Schmitt-Hoffner et al., 2021) | Does FOXR2 expression identify sympathetic neuroblastoma patients with unfavorable outcomes? | Yes | Yes | |

| ID | Claim | Citation | Question | Claim Answer | Paper Answer | Answer Note |
|---|---|---|---|---|---|---|
| 9 | FOXR2 activation via enhancer hijacking is a defining genetic hallmark of central nervous system (CNS) neuroblastoma. | (Liu et al., 2022);(Schmitt-Hoffner et al., 2021) | Is FOXR2 activation via enhancer hijacking a defining hallmark of CNS neuroblastoma? | Yes | Yes | |
| 10 | Central nervous system (CNS) neuroblastoma is a distinct molecular entity. | (Liu et al., 2022);(Schmitt-Hoffner et al., 2021) | Is CNS neuroblastoma a distinct molecular entity? | Yes | Yes | |
| 11 | In CNS neuroblastoma tumors, FOXR2 specifically targets LHX6+/DLX+ neural lineages derived from the medial ganglionic eminence. | (Jessa et al., 2024) | Does FOXR2 target LHX6+/DLX+ medial ganglionic eminence neural lineages in CNS neuroblastoma? | Yes | Yes | |
| 12 | In LHX6+/DLX+ neural lineages derived from the medial ganglionic eminence, FOXR2 activates ETS transcriptional networks to drive oncogenesis. | (Jessa et al., 2024) | Does FOXR2 activate ETS transcriptional networks in LHX6+/DLX+ medial ganglionic eminence lineages to drive oncogenesis? | Yes | Yes | |
| 13 | Activation of ETS transcriptional networks by FOXR2 promotes cellular proliferation. | (Yang et al., 2025) | Does FOXR2-driven activation of ETS transcriptional networks promote cellular proliferation? | Yes | Yes | |
| 14 | Activation of ETS transcriptional networks by FOXR2 influences mitogen-activated protein kinase (MAPK) signaling. | (Yang et al., 2025) | Does FOXR2 activation of ETS transcriptional networks influence MAPK signaling? | Yes | Idk | Passage doesn't state ETS activation causes MAPK influence |
| 15 | FOXR2 directly binds to key transcription factors that coordinate the initiation of gliogenesis. | (Jessa et al., 2024) | Does FOXR2 directly bind key transcription factors that coordinate the initiation of gliogenesis? | Yes | Yes | |
| 16 | FOXR2 binding to key transcription factors that coordinate the initiation of gliogenesis suggests a role for FOXR2 in disrupting normal neural differentiation during tumor development. | (Jessa et al., 2024) | Does FOXR2 binding to gliogenesis-initiation transcription factors suggest it disrupts neural differentiation during tumor development? | Yes | Yes | |
